# Modeling IP_3_ induced Ca^2+^ signaling based on its interspike interval statistics

**DOI:** 10.1101/2022.12.20.521161

**Authors:** Victor Nicolai Friedhoff, Martin Falcke

## Abstract

Inositol 1,4,5-trisphosphate (IP_3_) induced Ca^2+^ signaling is a second messenger system used by almost all eukaryotic cells. Recent research identified 8 general properties of Ca^2+^ spiking common to all cell types investigated and demonstrated randomness of Ca^2+^ signaling on all structural levels. We suggest a theory of Ca^2+^ spiking starting from the random behaviour of IP_3_ receptor channel clusters mediating the release of Ca^2+^ from the endoplasmic reticulum. Spike generation begins after the absolute refractory period of the previous spike. According to its hierarchical spreading from initiating channel openings to cell level, we describe it as a first passage process from none to all clusters open while the cell recovers from the inhibition which terminated the previous spike. Our theory reproduces quantitatively all general properties for different IP_3_ pathways including the exponential stimulation response relation of the average interspike interval (ISI) T_av_ and its robustness properties, random spike timing with a linear moment relation between T_av_ and the ISI standard deviation and its robustness properties, sensitive dependency of T_av_ on diffusion properties, and non-oscillatory local dynamics. We explain large cell variability of T_av_ observed in experiments by variability of channel cluster coupling by Ca^2+^ induced Ca^2+^ release, the number of clusters and IP_3_ pathway components expression levels. We predict the relation between puff probability and agonist concentration, and [IP_3_] and agonist concentration. Differences of spike behaviour between cell types and stimulating agonists are explained by the different types of negative feedback terminating spikes. In summary, the hierarchical random character of spike generation explains all of the identified general properties.

## 1 Introduction

The IP_3_ induced Ca^2+^ signaling pathway translates extracellular signals in the form of plasma membrane receptor agonist concentrations into intracellular responses by increasing the cytosolic Ca^2+^ concentration in a stimulus dependent pattern [5, 75, 77, 25]. The concentration increase can be caused either by Ca^2+^ entry from the extracellular medium through plasma membrane channels, or by Ca^2+^ release from intracellular storage compartments. In the following, we will focus on IP_3_-induced Ca^2+^ release from the endoplasmic reticulum (ER), which is the predominant Ca^2+^ release mechanism in many cell types [74]. IP_3_ sensitizes Ca^2+^ channels (IP_3_Rs) on the ER membrane for Ca^2+^ binding, such that Ca^2+^ released from the ER through one channel increases the open probability of neighboring channels (Fig. 1) [7, 4]. This positive feedback of Ca^2+^ on its own release is called Ca^2+^-induced-Ca^2+^-release (CICR). The released Ca^2+^ is removed from the cytosol either by sarco-endoplasmic reticulum Ca^2+^ ATPases (SERCAs) into the ER or by plasma membrane Ca^2+^ ATPases into extracellular space.

**Figure 1:**
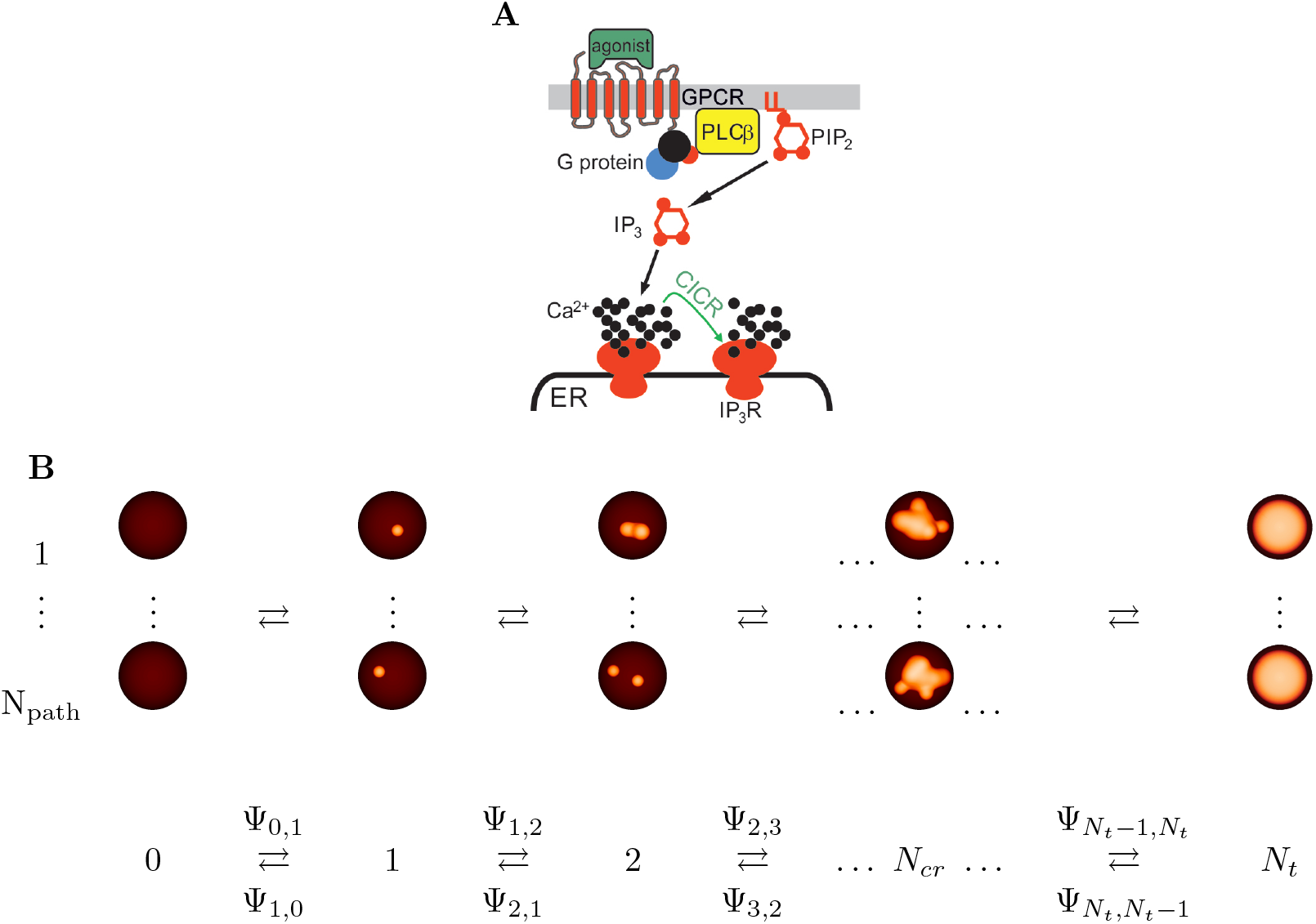
**(A)** The Inositol 1,4,5-trisphosphate (IP_3_) pathway. Binding of agonist to a G-protein coupled receptor (GPCR) activates Phospholipase C (PLC) which produces IP_3_ from Phosphatidylinositol-4,5-bisphosphate (PIP_2_). IP_3_ sensitizes IP_3_ receptor channels (IP_3_Rs) in the membrane of the endoplasmic reticulum (ER) for binding of Ca^2+^, such that Ca^2+^ released from the ER through one channel increases the open probability of neighboring channels by Ca^2+^-induced-Ca^2+^-release (CICR). **(B)** (Top) Open clusters are visualized as small orange spheres. A spike occurs when (almost) all *N*_*t*_ clusters are open. There are N_path_ paths of cluster openings and closing from 0 to *N*_*t*_ open clusters. (Bottom) Averaging over all paths leads to a state scheme indexed by the number of open clusters.

IP_3_Rs are spatially organized into clusters with a variable number of channels from 1 up to about 15. This cluster definition including single channels anticipates that the theory we will formulate is able to account also for channel populations which might not form clusters as suggested by Lock et al. [44]. Clusters are scattered across the ER membrane with reported distances of 1 to 7 µm [10, 67, 73, 71, 40]. CICR and Ca^2+^ diffusion couple the state dynamics of the channels. The coupling between channels ina cluster is much stronger than the coupling between adjacent clusters [76]. The structural hierarchy of IP_3_R from the single channel to clusters and cellular cluster arrays is also reflected by the dynamic responses of the intracellular Ca^2+^ concentration as revealed through fluorescence microscopy and simulations [10, 87, 46, 27, 66]. Random openings of single IP_3_R (blips) may trigger collective openings of IP_3_R within a cluster (puffs), while Ca^2+^ diffusing from a puff site can then activate neighbouring clusters, eventually leading to a global, i.e., cell wide, Ca^2+^ spike [46, 47, 40, 27, 66]. The timing of spikes is random [64, 54, 24, 21, 66, 80, 13, 15, 81, 56, 52, 39]. The typical timescale of spiking is the (temporal) average T_av_ of the interspike interval (ISI). Interestingly, that timescale cannot be found in long sequences of puffs from single isolated puff sites [79], i.e. it is an emergent property of the cellular dynamics. Repetitive sequences of Ca^2+^ spikes encode information that is used to regulate many processes in various cell types [5, 60, 42, 80].

At very strong stimulation, many Ca^2+^ signaling pathways exhibit a raised Ca^2+^ concentration of much longer duration than spikes, which may oscillate [3, 51], burst [33, 34] or be rather constant [50, 11, 38]. Typically, the amplitude of these oscillations is smaller than the spike amplitude.

Ca^2+^ exerts also negative feedback on the channel open probability, which acts on a slower timescale than the positive feedback, and has a higher half maximum value than CICR [87, 10, 47, 53, 35, 32, 79, 84, 45]. This Ca^2+^-dependent negative feedback helps terminate puffs, and therefore the puff probability immediately after a puff is smaller than the stationary value, but typically not 0. Channel clusters recover within a few seconds to the stationary puff probability in absence of other negative feedbacks [87, 10, 47, 53, 31, 35, 79, 45, 85].

IP_3_ is produced by Phospholipase C (PLC) upon stimulation of plasma membrane receptors. It sensitizes the IP_3_R for Ca^2+^ binding and thus signals from the plasma membrane to the puff probability [4]. Ca^2+^-mobilizing signals are generated by stimuli acting through a large variety of cell-surface receptors, including G-protein and Tyrosine kinases-linked receptors (Fig. 1) [6]. Stimulation of these receptors activates other pathways in addition to PLC, with plasma membrane receptor specific feedbacks to IP_3_ production and Ca^2+^ release [74, 6, 8, 1, 25, 55, 1]. They also affect the negative feedback terminating release spikes. Recovery from this negative feedback causes an *absolute* refractory period T_min_ as part of the interspike intervals (ISIs) lasting tens of seconds [86, 57, 80]. Hence, the negative feedback that determines the timescale of interspike intervals is different from the feedback contributing to interpuff intervals and requires global (whole cell) release events.

Recent experimental studies identified general properties of IP_3_-induced Ca^2+^ signaling - general in the sense that they apply to all cell types which have been investigated. We list these properties in the next section. Most of them are not captured by current theory of IP_3_-induced Ca^2+^ signaling. With this study, we would like to advance theory towards capturing these basic properties, and providing understanding of the stimulation response relation, cell variability and at the same time robustness of specific properties.

## 2 Results

### 2.1 Experimental results defining the mathematical theory

Identification of the mathematical structure to which IP_3_-induced Ca^2+^ dynamics corresponds requires to start from the basic observations applying to all cells. Some of these observations vary quantitatively between individual cells, others do not. We denote by cell variability differences of properties between individual cells of the same type and under the same conditions. We will denote properties which are the same in a quantitative sense for all individual cells of the same type stimulated with the same agonist as cell type specific and agonist specific. The average interspike interval 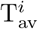 and its standard deviation *σ*^*i*^ are calculated as temporal average (temporal SD) over the ISIs of a spike sequence of an individual cell indexed by *i*. We develop our theory from these experimental observations:

1. Interpuff intervals (IPIs), puff amplitude and puff duration are random [53, 9, 31, 79, 20].

2. The time scale of Ca^2+^ spiking T_av_ does not appear in puff sequences of individual puff sites [79].

3. The sequence of interspike intervals is random [64, 54, 24, 21, 59, 80, 13, 15, 81, 56, 52].

4. The average interspike interval depends sensitively on the strength of spatial coupling between IP_3_R clusters [64, 63].

5. The standard deviation *σ*^*i*^ of interspike intervals obeys a linear moment relation to the average 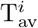:

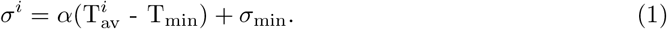

The slope *α, σ*_min_ and T_min_ are cell type and agonist specific and not subject to cell variability [64, 63, 54, 24, 21, 80, 13]. The slope *α* is robust against application of a variety of drugs and buffer addition [64, 63, 80]. *σ*_min_ is the standard deviation at saturating stimulation.

6. The average interspike interval 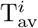 obeys

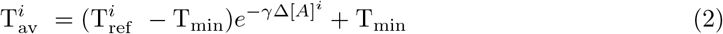

with the concentration of extracellular agonist 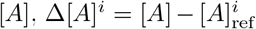, the concentration at the onset of spiking 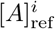, and the agonist sensitivity *γ*, which is cell type and agonist specific and not subject to cell variability. The exponential dependency has been observed for all pathways investigated [80].

7. Cell variability of the average interspike interval is large. It presents itself as large variability of the average interspike interval at the onset of spiking at low stimulation, and enters the stimulation response relation only via 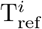 [64, 54, 24, 21, 59, 80].

8. Cellular release spikes are terminated by cell type and agonist specific negative feedback. That entails the puff probability immediately after a spike to be smaller than the asymptotic value after recovery from the negative feedback [57, 14, 47, 69].

Puffs and single channel openings (blips) are the noisy microscopic, elemental events, the sum of which constitutes the macroscopic cellular Ca^2+^ transient. Observation 1 provides a scale of the noise generated by the elemental events. Interpuff intervals are about one tenth of interspike intervals and local concentrations in the vicinity of puffs reach the level of global concentrations during a spike. The ratio in size and timescale of macroscopic to microscopic processes for many cellular systems is 10^5^ or more, which is sufficient to provide deterministic mean field behavior on cell level averaging out the microscopic noise. In the case of Ca^2+^ dynamics, that ratio is orders of magnitude smaller and we should not expect noise to average out. Thus, we need a stochastic approach from the start. The reason for this prominent role of noise is that a single random channel opening releases hundreds of Ca^2+^ ions per millisecond, and thus the molecular random event is amplified by several orders of magnitude.

Puffs and blips are not only the elemental events of Ca^2+^ transients, but also the local dynamics of a reaction-diffusion system. We learn from observation 2 that the local dynamics are non-oscillatory and that IP_3_R clusters do not reflect the spiking dynamics. Observation 3 confirms our expectation that microscopic noise does not average out on cellular level of this reaction-diffusion system. Observation 4 prohibits the formulation of the theory in terms of spatial averages.

We can draw some information on the interspike interval distribution from observation 5. The average T_av_ and standard deviation *σ* are not independent. This strongly suggests that the process setting T_av_ sets also *σ*. If we dealt with a noisy deterministic oscillator, some of the parameters setting the noise (*σ*) would most likely be different from the parameters setting the period, thus preventing a universal relation between average and standard deviation [65, 82].

The stimulation response relation is exponential (observation 6). Experiments established the exponential dependency on the agonist concentration not only on the basis of its excellent fits to measured relations, but also on the basis of a differential equation dT_av_ /d[A]=-*γ*T_av_ derived from stimulation-step experiments [80]. The experiments also showed that *γ* does not depend on the agonist concentration [A]. Since this relation is so solidly confirmed, we will use it to parameterize our theory. The agonist sensitivity *γ* is cell type and agonist specific. Surprisingly, the stimulation response relation does not show the canonical behavior close to any bifurcation generating limit cycles, indicating that the onset of spiking does not represent such a bifurcation in mathematical terms.

Cell variability of the average ISI T_av_ is a basic phenomenon, and is large (observation 7). That poses the question of the meaning of frequency in individual cells. Surprisingly, cell variability enters the stimulation response relation in this very defined way as the pre-factor of the exponential. The consequence for modeling is that models need to provide an explanation for cell variability and the robustness properties of the stimulation response relation.

The negative feedback terminating release spikes mentioned in observation 8 develops once a spike has started, i.e., when the Ca^2+^ concentration is high across the whole cell. It is a global process. It acts on all clusters in the cell simultaneously, and recovery from it also proceeds synchronously for all clusters. Therefore, we can assume spike termination and processes related to this feedback to be much less affected by noise than spike initiation. Hence, it obeys spatially averaged deterministic dynamics.

### 2.2 Defining the theory on the basis of experimental results

We start with formulating a theory for spike generation in a spike sequence. All channels and clusters are closed at the end of a spike [47], because the global negative feedback which terminated the spike has decreased the open probability substantially (observation 8) and started the absolute refractory period T_*min*_. When T_*min*_ has passed, channels and clusters start again to open and close sequentially till the next spike terminates. Due to recovery from negative feedback, all cluster opening probabilities increase proportional to 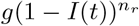 from 0 right after T_*min*_ (*t*=0, *I*=1) to their asymptotic value *g* (*t*=0, *I*=0). The variable *I* describes the negative feedback which terminated the previous spike. Its value increases during the spike, and we defined it such that the value is 1 at the time of spike termination.

The dynamics of the clusters are stochastic (observation 1). Each opening cluster entails a sphere of increased Ca^2+^ concentration around it. We indicate that by the orange spheres in the red round cells in scheme Fig. 1B. The local rise in Ca^2+^ increases the opening probabilities of the open cluster’s neighbours due to CICR (observation 4). A spike occurs, when (almost) all *N*_*t*_ clusters are open. This state is reached via one of many possible paths of cluster openings from 0 to *N*_*t*_ open clusters. Averaging over all N_path_ paths radically simplifies the system into a state scheme defined by the number of open clusters only [78], Fig. 1. Now, the ISI calculation corresponds to calculating the time it takes to get from 0 to *N*_*t*_ open clusters, which is called a first passage time [82]. The first passage time distribution corresponds to the ISI distribution for stationary spike trains.

The transition probabilities from *k* to *k*-1 open clusters are determined by the probability that one out of *k* open clusters closes with rate *δ*:

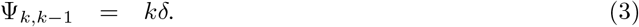

The probability for the first opening of a cluster is the single cluster puff probability times the number of closed clusters,

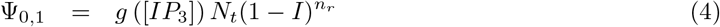

The inhibitory variable decreases towards 0 after spike termination with the rate *λ*, i.e.

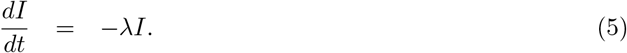

with the solution *I*(*t*) = *e*^*−λt*^. The puff probability depends on the *n*_*r*_*th* power of 1-*I*.

The transition probabilities from *k* to *k*+1 open clusters pick up Ca^2+^ induced Ca^2+^ release (CICR). [Ca^2+^] increases with the number of open clusters - either in the whole cell or at least in the vicinity of open clusters. We assume this concentration increase to be proportional to the number of open clusters *k*. Each open cluster increases [Ca^2+^] by *s*_*p*_*c*_*r*_*k* with *c*_*r*_ denoting the resting concentration. That rise of concentration increases the channel opening rate due to CICR. Thus, CICR introduces an increase of the transition probabilities from *k* to *k*+1 with increasing *k*. Two factors determine how quickly Ψ_*k,k*+1_ increases. The first one is the dependency of the channel open probability on [Ca^2+^]. According to a variety of studies, the open probability increases like 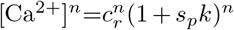 [7, 18, 48, 30, 62, 26], with reported values for *n* from 1.0 to 4.0 (1.0-2.7 [30], 2.7 [7], 1.6 our fit to data from [62], up to 4.0 [48]). The second factor derives from the picture of spike generation as wave nucleation. The number of closed clusters neighbored to the expanding wave increases like the surface of the volume engulfed by the wave, i.e. like *k*^2*/*3^. These clusters contribute most to the [Ca^2+^]-mediated increase in open probability. Combining this with the channel open probability dependency on [Ca^2+^] suggests *n*=1.7-4.7. So, while we are aware that our choice *n*=3 may not apply to all cell types or situations, we think it covers many situations.

The constant *s*_*p*_ accounts for the strength of spatial coupling and depends on many factors. The level of Ca^2+^ in the ER determines the release current and thus the concentration amplitude of a puff [2]. The very geometry of the cell-wide cluster array sets the distances and thus the [Ca^2+^] increase due to diffusion. Buffers set the effective diffusion coefficient. This non-exhaustive list illustrates that this CICR-based coupling is a likely candidate for the cause of cell variability. At the same time, the convergence of so many biological parameters on a single parameter of the dynamics is one reason for the robustness and universal properties of Ca^2+^ spiking. Including all factors, we obtain the transition rate from *k* to *k* + 1

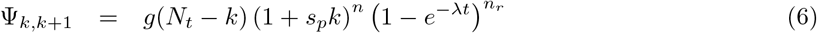

comprising the factors (asymptotic single cluster puff probability *g*)×(number of closed clusters *N*_*t*_ − *k*)×CICR×(recovery from negative feedback of previous spike). We subsumed the factor 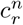 in *g* and have chosen *n*=3 in this study.

The average time scale of transitions between the states in the linear chain in scheme Fig. 1 is set by the average time scale of individual cluster interpuff intervals (puff duration) divided by the number of closed (open) clusters. Already the single cluster puff rate is much faster than the time scale of interspike intervals and thus Ψ_0,1_ implements observation 2.

The dynamics of the state probabilities is given by the Master Equation

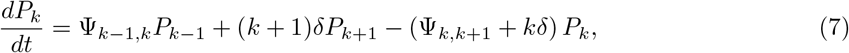

with *k* = 0, …, *N*_*t*_ *−* 1. Its Laplace transform defines the difference equation (S5) for the Laplace transforms of the state probabilities, which we solved analytically by Eq. (S6) [29]. The moments of the ISI distribution are determined by Eqs. (S7, S8). We also simulated trajectories and compare their outcome to the analytical calculations in Fig. (S2).

We have introduced our theoretical approach with clusters as stochastic elements and Ca^2+^ concentration profiles around open clusters. It would have the same structure if we assumed an array of single channels to generate a spike, as the observations by Lock et al. [44] suggest. The first passage formulation would also apply if we considered very small cells or very fast diffusion such that the concentration rise caused by open clusters is spatially homogeneous. We could even think of mixed forms, with clusters starting the release and more diffuse single channels joining later, by specifying the Ψ_*k,k±*1_ accordingly.

Summarizing this section, we established the mathematical structure to which we assume IP_3_-induced Ca^2+^ spiking to correspond. It is a random walk in the state space of the IP_3_R cluster array of a cell with transition rates depending on [Ca^2+^] and [IP_3_]. We radically simplified the state space to just the number of open clusters.

### 2.3 Applying the theory

#### 2.3.1 From puffs to spikes

Puffs and blips are the elemental release events, and only they are observed at low stimulation [10, 87, 46] or at very weak spatial coupling [17, 16, 88]. The transition from this regime to spiking happens when rising stimulation increases the puff probability. Fig. 2 illustrates the transition from puffs to spikes. It shows the average first passage time from all clusters closed at the end of the absolute refractory period T_min_ to *N* open clusters (*N* ≤ *N*_*t*_) and relates it to the single cluster puff probability *g*. Considering events of a given amplitude, i.e. a given *N*, all of them become more frequent with increasing puff probability.

**Figure 2:**
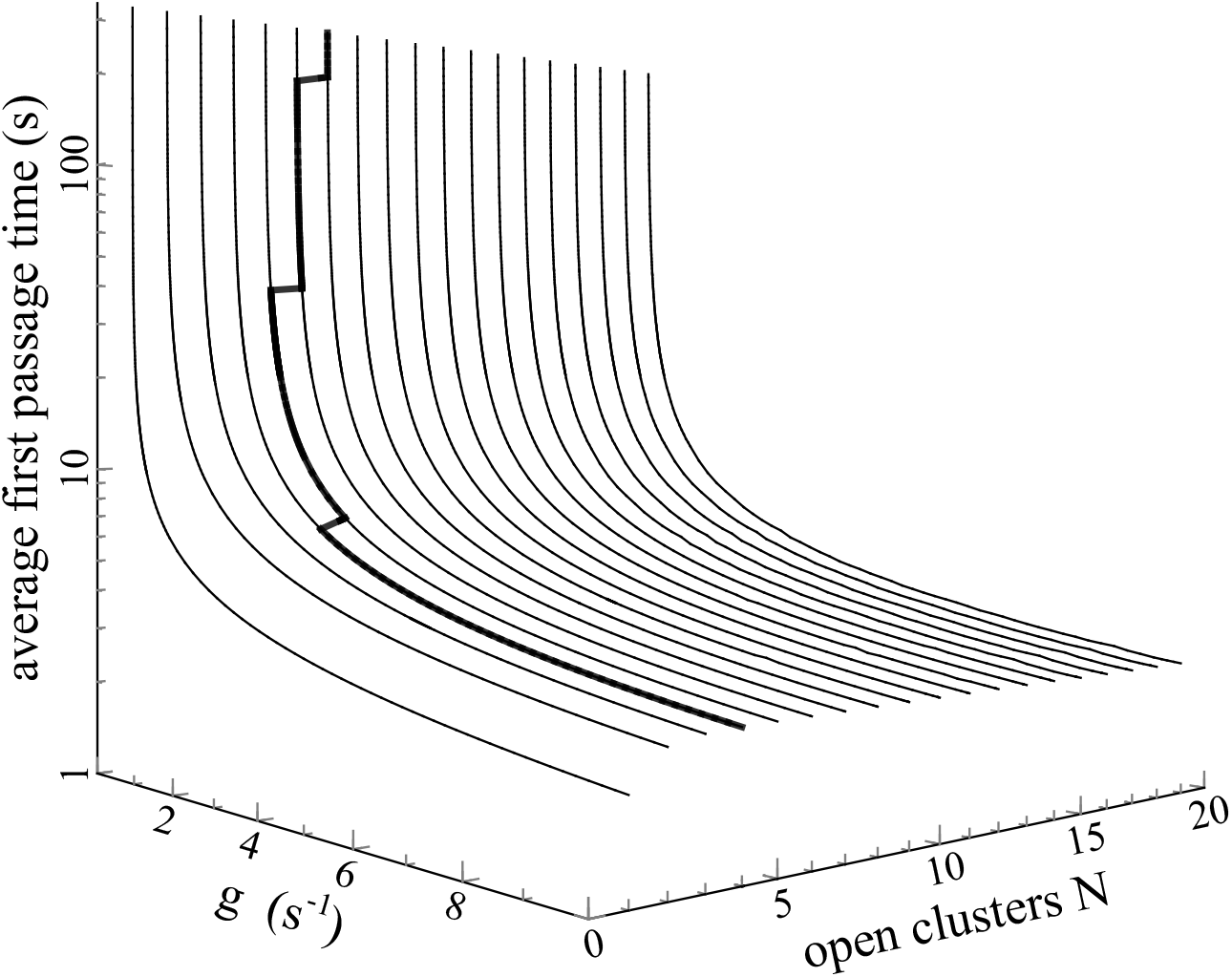
Average first passage time (FPT) from 0 to N open clusters in dependence on the single cluster puff probability *g*. The value of *g* is the puff probability after complete recovery from the previous spike. The parameters used are *λ* = 0.001443 *s*^*−*1^, *s*_*p*_ = 1.7, *N*_*t*_ = 30. Saturation of the average FPT in dependence on *N* starts at *N* = *N*_*cr*_(*g*), which is marked by the thick line.

Single cluster openings occur after about 10 s at very low values of *g* (with the parameter values in Fig. 2), but it takes a hundred or more seconds till 2 clusters are open at the same time and even longer for 3 open clusters or a spike. As a time course, we would see many puffs before a larger event happens. At large values of *g*, puffs are still a few times more frequent than events involving more clusters. But the time till large *N* -values are reached decreased substantially and spikes happen soon after the absolute refractory period.

Spike generation becomes more tangible by following the average first passage time with increasing *N* at a given large puff probability in Fig. 2. The first passage time saturates at a certain number of open clusters *N*_*cr*_. Once this value *N*_*cr*_ is reached by the random cluster opening, almost all clusters open in next to no time due to the strong positive feedback of CICR to the cluster opening rate. The average interspike interval T_av_ is essentially equal to the average first passage time from 0 to *N*_*cr*_ open clusters. The value of *N*_*cr*_ depends on the single cluster puff probability *g* and on the parameters setting it like [IP_3_], the extracellular agonist concentration, the molecular properties of IP_3_Rs and the strength of spatial cluster coupling by CICR. The existence of *N*_*cr*_ ≤ *N*_*t*_ corresponds to the experimental observation in a variety of cell types that opening of a few clusters starts a global spike with almost certainty [58, 9, 12, 27] (see also here ^1^).

#### 2.3.2 The dependency of the average ISI on cluster-cluster coupling and cluster number

We explained above that the many factors setting the strength of spatial coupling between IP_3_R clusters converge on the parameter *s*_*p*_. Fig. 3 illustrates the relation between T_av_ and *s*_*p*_. We see that the average ISI depends on the strength of spatial coupling (observation 4) described by parameter *s*_*p*_. The dependency of T_av_ on Ca^2+^ diffusion properties has been investigated by Skupin et al. [64] by measuring T_av_ before and after addition of the Ca^2+^ buffers BAPTA or EGTA to spiking cells. They substantially reduce the concentration of free Ca^2+^ in the vicinity of open channels corresponding to a decrease of *s*_*p*_ [76]. Only a part of cells resumed spiking after buffer addition and mostly those with small T_av_ before buffer was added. The average ISI of those continuing to spike increased by a factor of 1.7 on average. The examples in Fig. 3 cover this range of T_av_-change. T_av_ is most sensitive to *s*_*p*_ changes in the range of small *s*_*p*_ values. Cells in that range most likely corresponds to the fraction of cells in the experiments which did not resume spiking after buffer addition.

**Figure 3:**
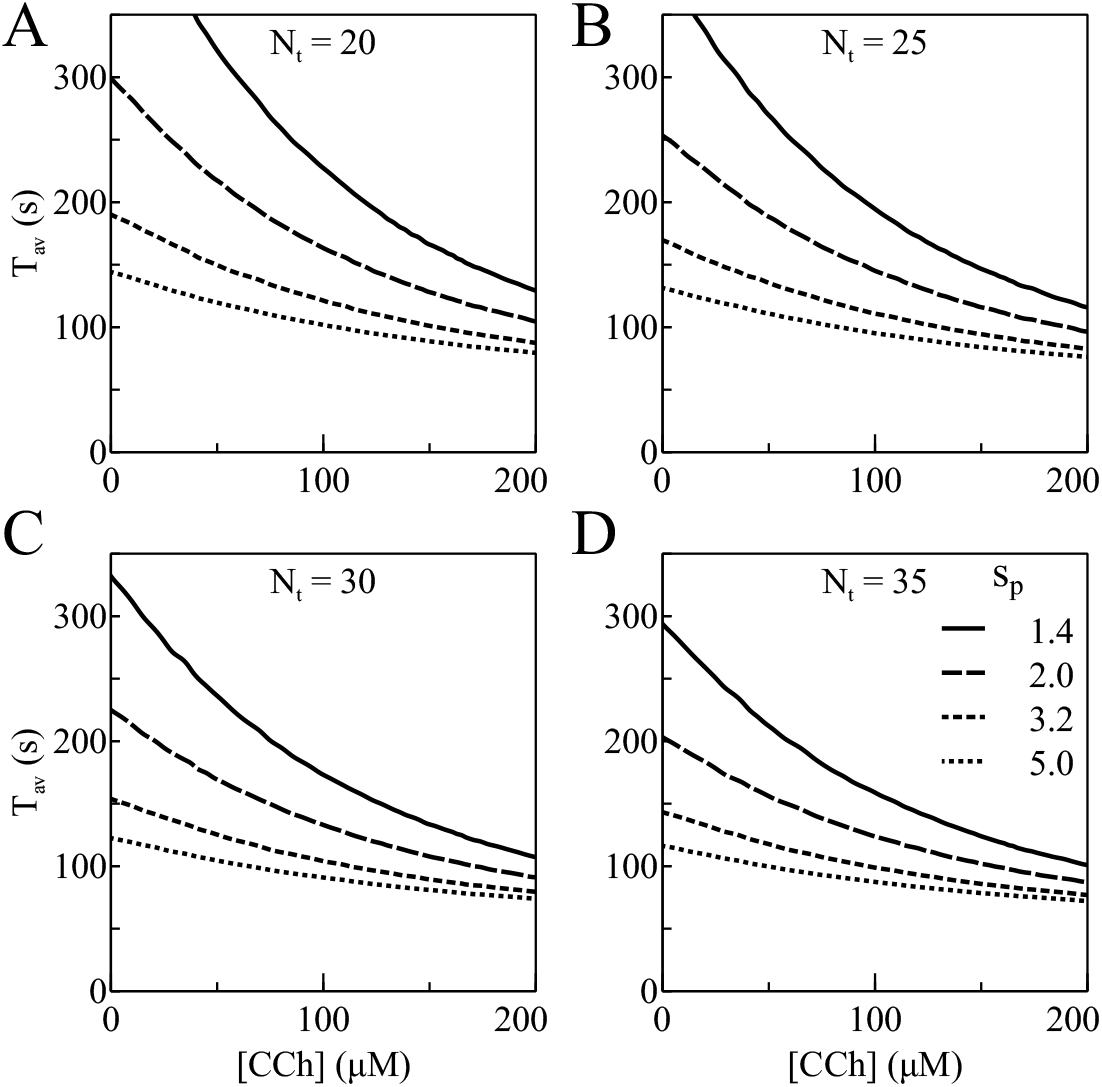
The average interspike interval is affected by the strength of spatial coupling *s*_*p*_ and the number of clusters in the cell *N*_*t*_. The four panels show T_av_([CCH]) for HEK cell parameters for four *N*_*t*_ values and four *s*_*p*_ values. The smaller the value of *s*_*p*_ - and therefore the weaker the spatial coupling - the larger is the average ISI. Note that T_av_ responds more sensitive to a change of *s*_*p*_ from 2.0 to 1.4 than from 5.0 to 3.2. The average ISI decreases with increasing *N*_*t*_ as a comparison across panels for equal *s*_*p*_-values shows. Legend in (D) applies to all panels.

The total cluster number in the cell *N*_*t*_ is the other geometrical parameter affecting the average ISI. T_av_ decreases with increasing cluster number *N*_*t*_ (Fig. 3). That is one of the fundamental differences to models based on mean field dynamics exhibiting a dependency of rates on concentrations. With a dependency of rates on numbers, increasing the cell size while keeping the IP_3_R concentration constant decreases T_av_ (see [27, 66] for simulations).

#### 2.3.3 The slope of the moment relation

The moment relation Eq. 1 has been confirmed for all cell types and pathways investigated so far. Its linearity is an important constraint to mathematical models, and the value of its slope *α* indicates the timescale of recovery from the negative feedback terminating release spikes [78]. Hence, it contains information on the pathways co-activated with PLC by cell stimulation (see Fig. 1).

We identify *α* with the coefficient of variation (CV) of the underlying stochastic process, given by 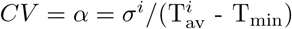. The range of possible values for *α* is therefore between 0 for a deterministic spiking process and 1 for a homogeneous Poisson process. The experimentally determined value of HEK cells for a variety of CCh concentrations and co-application of other drugs varied in a rather small range between 0.20 and 0.28 [80]. We consider *α* as robust and not subject to cell variability on that basis. Consequently, the main determinant of the value of *α* can only be other parameters not subject to cell variability. Indeed, the rate of recovery from negative feedback *λ* and the exponent *n*_*r*_ turned out to be the main determinants [78].

We used *α* and the range of observed T_av_-values as fit criteria to determine the value of *λ* and *n*_*r*_ applying to the different cell types and agonists (Table 1). The moment relations resulting from these fits are shown in Fig. 4 and reproduce the measured *α*-values. Note, the slower the recovery from negative feedback (the smaller *λ*), the smaller the value of *α* [29]. The *α*-values for Astrocytes, HEK cells and Hepatocytes stimulated with Phenylephrine or Vasopressin could all be fit with the exponent *n*_*r*_=1.0 in the recovery factor (Table 1).

**Table 1.** Parameter values varying between pathways and cell types. Parameter values identical for all cell types are: *δ*=6.93 s^*−*1^ [20, 19], *n*=3 [7, 18, 48, 30, 62, 26], [IP_3_]_0_=0.01 ms UV flash, *k*_*p*_=0.016 ms^*−*1^ [20], *g*_0_= 5.0 s^*−*1^ [79], *n*_*r*_=1. Parameters describing cell variability are *s*_*p*_=1.0-5.0 and *N*_*t*_=15-45. Their value ranges have been chosen to match observed T_av_ ranges. We consider *g*_0_ as cell type and agonist specific (see text). Since we do not have measured values for the different pathways, we fixed it to a value allowing to describe the observations in all 4 pathways. The same applies to [IP_3_]_0_.

| Pathway | $\lambda$ | $T_{\min}$ |
| --- | --- | --- |
| HEK cells, CCh | $1.443 \cdot 10^{-3} \text{ s}^{-1}$ | 57 s |
| Hepatocytes, Pe | $1.154 \cdot 10^{-2} \text{ s}^{-1}$ | 61 s |
| Hepatocytes, Vp | $6.0 \cdot 10^{-4} \text{ s}^{-1}$ | 44 s |
| Salivary gland, 5-HT | $1.154 \cdot 10^{-3} \text{ s}^{-1}$ | 16 s |
| Astrocytes, spontaneous | $4.0 \cdot 10^{-1} \text{ s}^{-1}$ | 54 s |

**Figure 4:**
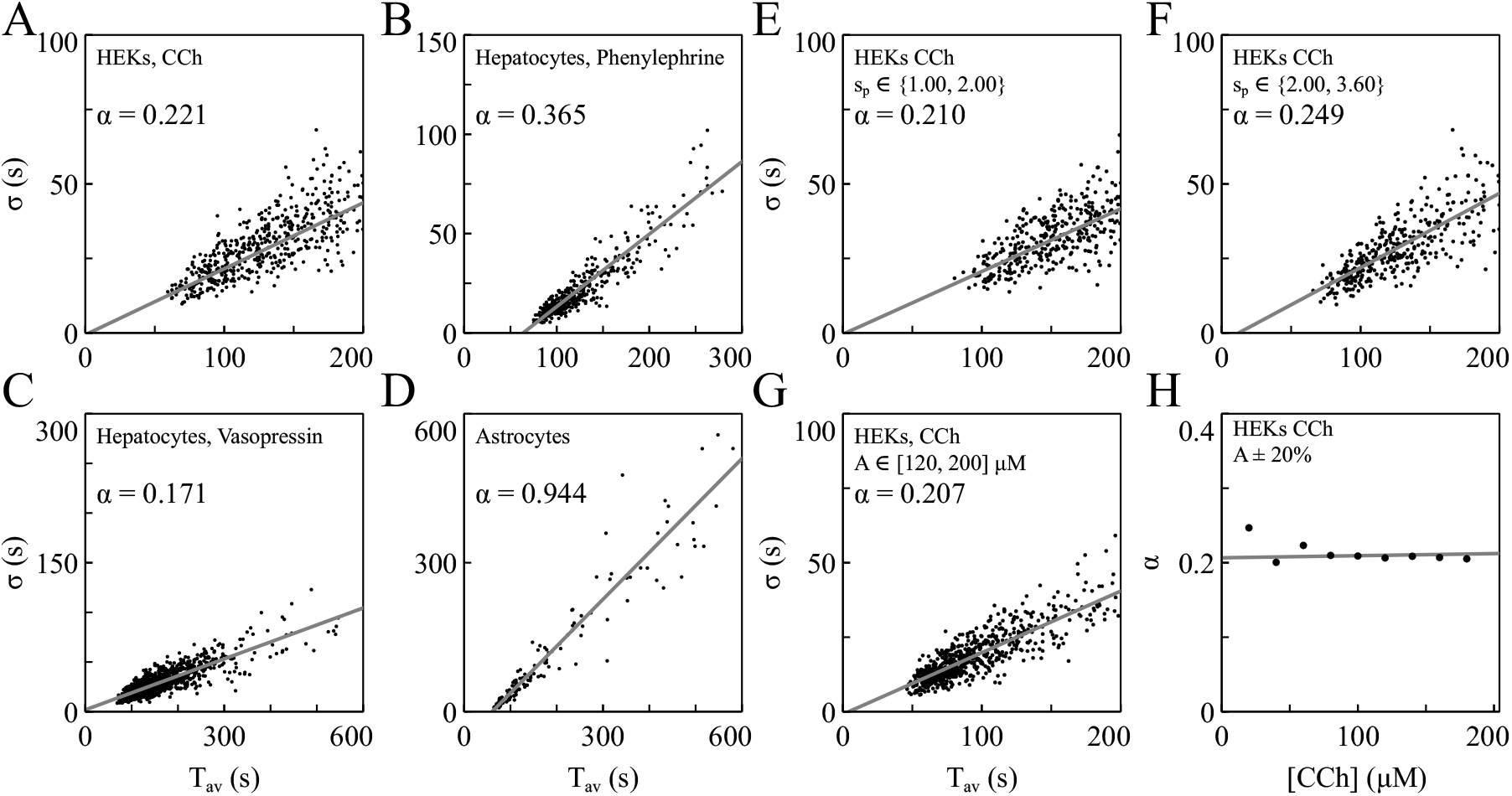
Moment relation (1) between the average ISI T_av_ and the ISI standard deviation *σ* for four pathways. Parameter values are listed in Table 1. **(A)** HEK cells stimulated with [CCh] ∈[20, 100] *µ*M. The measured range of *α* is 0.20 to 0.28 [80]. **(B)** Hepatocytes stimulated with Phenylephrine. The measured value of *α* is 0.37 [80]. **(C)** Hepatocytes stimulated with Vasopressin. The measured value of *α* is 0.17 [80]. **(D)** Spontaneously spiking Astrocytes. The measured range of *α* is 0.94-1.01 [63]. Analogous to the analysis of experimental data in [80, 63], averages over 12 consecutive ISIs sampled from simulations were used for each data point. Data points differ in *N*_*t*_ and *s*_*p*_. **(E, F)** *σ*-T_av_ plot for the HEK cells, *α* has been determined with different *s*_*p*_ values while all *N*_*t*_ are included and [CCh] as in panel (A). *α* varies slightly but stays in the experimentally measured range, showing that it is a robust property of the system w.r.t to a change of factors affecting *s*_*p*_. **(G)** *σ*-T_av_ plot for HEK cells with the same parameters as in (A) apart from the stimulation range now [CCh] ∈ [120, 200] *µ*M. **(H)** *α* changes only slightly with agonist concentration and stays within the measured range. Data points for a given [CCh]-value were determined with *σ* and T_av_ values within [CCh] ± 20%.

The value of *α* is surprisingly robust. Addition of buffer, co-application of drugs with stimulation, which changed IP_3_ production, IP_3_ sensitivity of the IP_3_R or ER Ca^2+^ uptake, and varying stimulation all changed T_av_ but not *α* [64, 63, 80]. Modeling the robustness of *α* requires first to remind us of likely properties of cell variability. We discussed before that all the factors setting the strength of spatial coupling *s*_*p*_ and the number of clusters *N*_*t*_ are likely candidates for causing cell variability. Expression levels of the components of the IP_3_ pathway like the plasma membrane receptor or Phospholipase C most likely also vary between individual cells causing cell variability in agonist sensitivity and *g*([A]). Indeed, it is a common experience that individual cells start to spike at different stimulation strength. That means in terms of the model that the value of 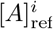 in Eq. 2 varies between cells (as index *i* indicates).

The moment relation is completely determined by T_av_(g) and its dependency on *s*_*p*_ and *N*_*t*_. Hence, components of the IP_3_ pathway affect the moment relation only via *g* and it suffices to demonstrate its robustness against changes of *g*. We vary [CCh], *s*_*p*_ and *N*_*t*_ to change *g*.

Fig. 4 panels E-G show the moment relation with HEK cell parameters for ranges of parameter values describing cell variability. *N*_*t*_, *s*_*p*_ and 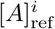 have been varied to obtain cell variability. Additionally we vary the ranges from which we sample *s*_*p*_ and [CCh]. Fig. 4H shows the value of *α* for the range of [CCh] used in experiments. The value of *α* stays always well within the measured range. These results demonstrate that our theory reproduces the robustness properties of *α*.

#### 2.3.4 The relation between puff probability and agonist concentration and the exponential stimulation response relation

We quantify the relation of the puff probability to the agonist concentration *g*([A]) by using the stimulation response relation T_av_([A]) (2) and T_av_(*g*) from Fig.2 to obtain T_av_(*g*)=T_av_([A]) relating *g* to [A] via the T_av_ values (see supplemental material Eqs. S12-S14). Rearranging that equation for *g* provides *g*([A]).

The dependencies of the puff probability *g* on agonist concentration *g*([A]) obtained with the measured stimulation response relations from ref. [80] are shown in Fig.5. They have been determined with typical values of *s*_*p*_ and *N*_*t*_ for each cell type. We call them for later use reference relations.

**Figure 5:**
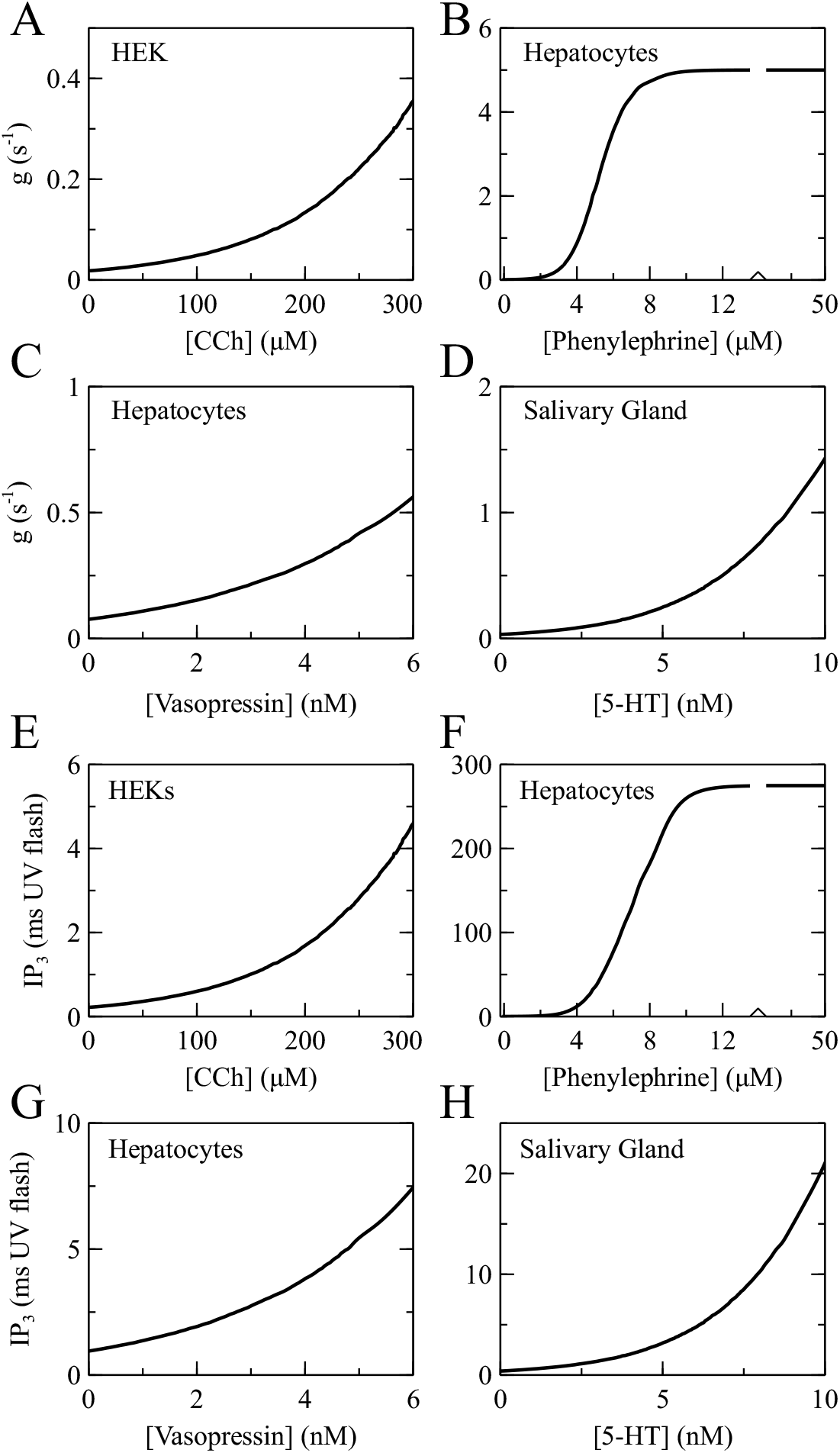
**(A-D)** Single cluster puff probability *g* vs agonist concentration for four pathways. The stimulation response relation Eq. 2 saturates for Hepatocytes stimulated with Phenylephrine, and so does *g*([Phenylephrine]). The stimulation response relation does not saturate with the other three pathways, and therefore neither reaches *g*([A]) saturation within the spiking regime. **(E-H)** IP_3_([A]) for four different pathways. The reference puff probability agonist relations were used for each pathway.

As expected, *g* increases with stimulation in all cases. We determine *g*([A]) for the agonist concentration range covered by the measured stimulation response relations. If these relations do not saturate within the range of measured agonist concentrations, neither does *g*([A]). That applies to HEK cells, salivary gland cells and Hepatocytes stimulated with Vasopressin in Fig. 5. Stimulating Hepatocytes with increasing concentration of Phenylephrine (Pe) leads to saturation of the average ISI at T_min_ at a few *µ*M. These cells still spike at 50 *µ*M, and the corresponding relation *g*([Pe]) saturates and has a sigmoidal shape.

We use *g*([A]) to learn more about the stimulation response relation, now. The population average of the experimentally measured stimulation response relation can be fit by a single exponential [80]. That strongly suggests that this relation for all individual cells obeys essentially the same exponential function and is not affected by cell variability [80]. Cell variability enters by the pre-factor of the exponential. The surprising aspects of this observation are that given all the obvious cell variability among the pathway components, the exponential relation is not violated, all cells have the same value of *γ* and cell variability enters so well-defined.

There is no indication or a priori reason to assume that the parameters causing cell variability are correlated in a way guaranteeing the exponential relation and the value of *γ*, like for instance that cells with small *N*_*t*_ have a very sensitive IP_3_ pathway to compensate for the small cluster number. Rather, the ability to fit the concentration response relations for a range of the parameter values describing cell variability with the same relation *g*([A]) and by varying the pre-factor of the exponential only is compatible with our ideas on cell variability. We use the reference relation *g*([A]) determined above to verify this ability of our theory.

We show stimulation response relations for four different pathways in Fig. 6. The full line in each panel corresponds to the measured relation in Thurley et al. [80]. It is exactly reproduced by definition of the reference *g*([A]). Other line styles show stimulation response relations with other values of cell variability parameters than the reference relation. The exponential relation with the measured agonist sensitivity *γ* is a very good approximation for the stimulation response relations of cells with different values of *s*_*p*_ and *N*_*t*_ than the reference relation in all four cases in Fig. 6. Variability in 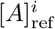does not cause deviations from the exponential dependency. It merely changes 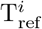 due to the properties of the exponential function in agreement with the experimental results. Hence, our theory is able to explain the robustness properties of the stimulation response relation observed by Thurley et al. [80] without assuming compensating correlations between the parameters of cell variability.

**Figure 6:**
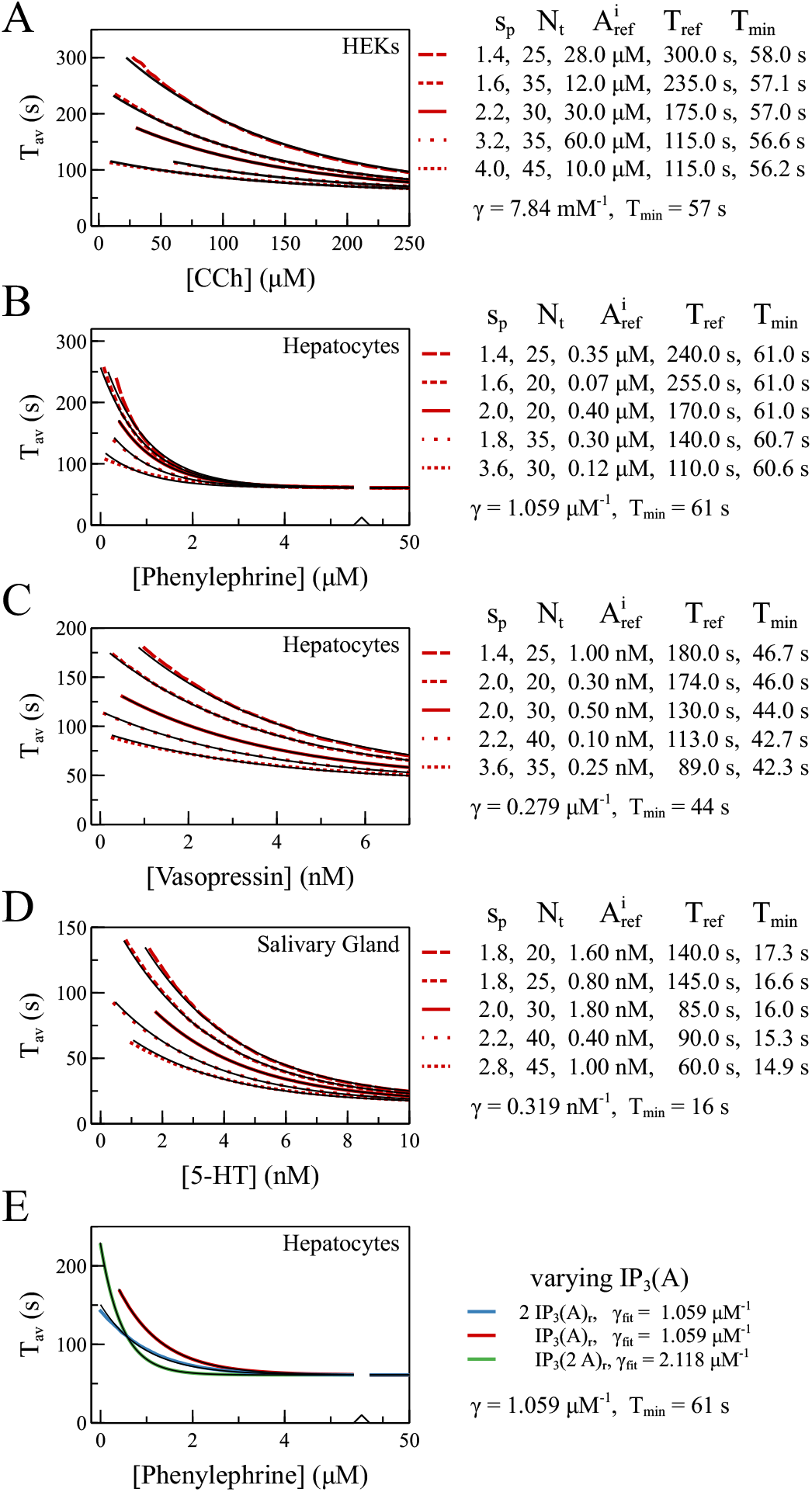
**(A)-(D)** T_av_([A]) for different pathways and different combinations of *N*_*t*_ and *s*_*p*_ are shown. Red lines are fits to exponential functions. Black lines are calculated from the stochastic process of spike generation and the reference relation g([A]) as T_av_(g([A]),*s*_*p*_,*N*_*t*_), i.e. we first calculate g from [A] (Fig. 5A-D) and then T_av_(g) (Fig. 2). The data shown by solid lines use the reference values for *s*_*p*_ and *N*_*t*_. They fit the experimental data from ref. [80] perfectly by definition. All other lines use different values of *s*_*p*_ and *N*_*t*_ but the same reference g([A]). Fits are exponential functions from observation 6 using the pathway specific value for *γ* for all lines, and T_ref_ and T_min_ as fit parameters. Hence, the curves differ in T_ref_ and by small amounts in T_min_. The quality of the fits illustrate that the stimulation response relations of a given pathway for different cell variability parameter values can be very well approximated by the same exponential and T_min_ values. That reproduces the robustness properties of the stimulation response relation. **(E)** We show T_av_(g([IP_3_]([A])),*s*_*p*_,*N*_*t*_), i.e. we first calculate [IP_3_] from [A], then g from [IP_3_] ((8)) and then T_av_(g). Additionally, we show T_av_(g(2[IP_3_]([A])),*s*_*p*_,*N*_*t*_) to show the effect a two-fold expression level of the IP_3_ pathway, and T_av_(g([IP_3_](2[A])), *s*_*p*_, *N*_*t*_) for doubled sensitivity. All calculations use the reference values for *s*_*p*_ and *N*_*t*_. Colored lines show exponential fits. Doubling the sensitivity of [IP_3_]([A]) doubles the value of *γ*.

While the exponential dependency on agonist in T_av_([A]) was an input to our theory, the robustness properties of T_av_([A]) were not. Therefore, we consider the finding that they are met by our theory as a confirmation of our choices.

#### 2.3.5 The relation between [IP_3_] and agonist concentration

Binding of agonist to a G-protein coupled receptor (GPCR) activates PLC, which produces IP_3_ diffusing to the IP_3_R and sensitizing the receptor for binding of activating Ca^2+^ (Fig. 1A). Knowledge on the concentration of the intracellular messenger IP_3_ was not required for the results presented so far, since we know the stimulation response relation of T_av_. Indeed, we need additional experimental information to be able to make statements on [IP_3_]. According to our approach starting from puff probability, we invoke the relation between [IP_3_] and puff probability measured by Dickinson et al. [20]. The expression

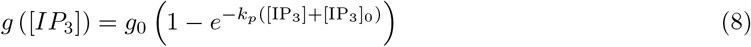

is a very good fit to the data in Dickinson et al. [20]. It is set up to [IP_3_] in units of UV flash duration used in experiments uncaging caged IP_3_. Dickinson et al. measured in SH-SY5Y cells with a maximum puff probability *g*_0_ of about 1 s^*−*1^. The measurements in Thurely et al. suggest it to be up to 5 times larger in HEK 293 cells [79]. Hence, we expect the maximum puff probability *g*_0_ to be specific to the agonist and cell type.

We can now determine the relation between [IP_3_] and agonist concentration, [IP_3_]([A]), by inverting (8) to provide [IP_3_](g) and using *g*([A]) for [IP_3_]([A])=[IP_3_](g([A])) (Fig. 5). It is an effective description of the behaviour of the GPCR and PLC controling IP_3_ production. The same remarks as for *g*([A]) with regard to the agonist concentration range and saturation behavior apply to [IP_3_]([A]). We see sigmoidal behavior for [IP_3_]([Phenylephrine]). This saturating type of relation is typical for the response of signaling pathways. The other three pathways do not saturate within the agonist concentration range available for the fit. However, their shape is compatible with sigmoidal functions saturating at larger concentrations maybe beyond the spiking in the over-stimulation regime. We used *g*([A]) to derive [IP_3_]([A]). That guarantees consistency with the relation between puff probability *g* and T_av_ as well as the stimulation response relation (2). This consistency in mathematical terms means

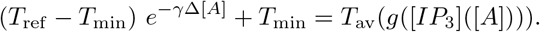

The red line in Fig. 6E shows this consistency for the reference case.

We expect cell-to-cell variability also in the expression level of the components of the pathway leading to IP_3_ production like e.g. the GPCR, G protein and PLC (see Fig. 1A). However, in difference to the moment relation and the stimulation response relation, we do not have experimental data verifying this assumption or quantifying the robustness properties. Therefore, we are limited to theoretical considerations. We expect the stimulation response relation to exhibit robustness with regard to changes of some properties of the IP_3_ pathway, but also to reflect the cell type and agonist specificity of *γ*, i.e. some properties of the IP_3_ pathway will affect *γ*.

The expression level of IP_3_ pathway components fixes the saturation value [IP_3_]_max_ of [IP_3_]([A]) (see Fig. 5F). Fig. 6E shows T_av_(*g*([IP_3_]([A]))) with doubled [IP_3_]_max_. It can be fit very well with the same exponential as the reference case. Changing the saturation value from 70% to 200% of its reference value affects mainly the pre-factor and the agonist sensitivity *γ* by ± 10% only (Fig. S5). Hence, the expression level of the pathway components affects the stimulation response relation in good agreement with its robustness properties in a large range of [IP_3_]_max_. Interestingly, T_av_(*g*([IP_3_]([A]))) was more robust against an increase of [IP_3_]_max_ than a decrease. We investigated whether values of the parameters determining [IP_3_]([A]) exist, providing more robustness against decreasing [IP_3_]_max_. Fig. S5 shows that this clearly is the case. Increasing either *k*_*p*_ or *g*_0_ extends the robust range towards small [IP_3_]_max_. However, a definite statement on the robustness properties requires more experimental research to position measured [IP_3_]([A]) in parameter space.

The sensitivity of the pathway is determined by molecular properties like the specific type of GPCR, Tyrosine kinase and PLC involved [4], which are not subject to cell variability. The sensitivity of [IP_3_]([A]) should therefore affect elements of our theory with the same robustness properties. That is what we observe when we double the sensitivity of [IP_3_]([A]) by use of [IP_3_](2[A]). It doubles the agonist sensitivity of the stimulation response relation *γ* according to our expectations (Fig. 6E).

Mathematically, *g*([IP_3_]([A])) transforms T_av_(g) into the stimulation response relation. Since *g*([IP_3_]) ((8)) and the stimulation response relation ((2)) are fixed by experimental results, the outcome of the stochastic process of spike generation T_av_(g) (Fig. 2) shapes [IP_3_]([A]). The result that [IP_3_]([A]) has a sigmoidal shape as we expect for a signaling pathway and is compatible with the robustness properties of the stimulation response relation are confirmations for the assumption that spike generation is a first passage process.

## 3 Discussion

Our theory describes the complete pathway from the agonist concentration to the Ca^2+^ spiking characteristics. Stimulation with agonist produces [IP_3_] according to the relations shown in Fig. 5. That entails a puff probability *g*([*IP*_3_]([*A*])) according to (8). The value of the puff probability finally entails an average interspike intervall according to the relation 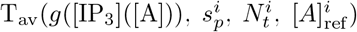 calculated from the stochastic process of spike generation with examples shown in Fig. 2. The parameters 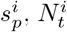 and 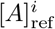 account for cell variability and specify individual cells. The complete pathway model is in agreement with all the general observations listed in section 2 and additionally with (8).

Cell variability of T_av_ is large. At the same time, it is an important source of information on the relations defining IP_3_ induced Ca^2+^ spiking quantitatively and their robustness properties. We established the moment relation and stimulation response relation by exploiting the cell variability found in experiments [64, 80]. In this study, we used the robustness properties revealed by the features not subject to cell variability as crucial information for verifying and parameterizing our theory. Cellular parameters with given robustness properties can only be related to parameters with equal or even more general robustness. That is a criterion for defining parameter relations. We confirmed it for the agonist sensitivity *γ* by showing that it does not depend on the cell variability parameters. We exploited it by fixing the recovery rate *λ* with the slope *α* of the moment relation.

Cell type and agonist specific properties enter the theory by 3 experimentally determined parameter values. The moment relation provides the slope *α* corresponding to the coefficient of variation (CV) of the stochastic part of the interspike interval T_av_ -T_min_. It fixes the recovery rate *λ*. The stimulation response relation contributes the value of the agonist sensitivity *γ* and the absolute refractory period T_min_. They determine the relation between single cluster puff rate and agonist concentration *g*([A]), and affect via (8) also [IP_3_]([A]).

We have chosen the parameters *s*_*p*_, *N*_*t*_ and [*A*]_ref_ to describe cell variability. The choice of *s*_*p*_ and *N*_*t*_ is obvious from the vast experimental literature showing puff sites in individual cells. The value ranges have been chosen to match the observed T_av_ ranges. The common experience that individual cells start spiking at individual agonist concentrations is picked up by cell specific [*A*]_ref_.

Our theory comprises 5 additional parameters which we kept constant in this study: *n, n*_*r*_, *δ, k*_*p*_, [IP_3_]_0_. We kept the cluster closing rate *δ* constant across all cell types and pathways, since recent studies showed that all IP_3_R isoforms produce similar puffs [43, 49]. *n, δ* and *k*_*p*_ are fixed by experimental results (see Table 1).

IP_3_ induced Ca^2+^ spiking generates spikes in a hierarchical process and exhibits randomness on all structural levels. Our theory starts from these observations. Spiking in general obeys the linear moment relation ((1)) and the exponential stimulation response relation ((2)), IP_3_ induced puffs obey (8), and so does our theory including the corresponding robustness characteristics. These relations are mechanistically unrelated observations without theory. Calculating T_av_(g) from the stochastic first passage process was sufficient to provide a quantitative picture connecting all three relations and predicting the relation between puff probability *g* and stimulation [A], *g*([A]), and [IP_3_] and stimulation [IP_3_]([A]).

### 3.1 Comparing modeling concepts

Early modeling concepts of intracellular Ca^2+^ signaling pioneered mathematical modeling in cellular physiology [22, 18, 72, 68, 26]. They derived mean field differential equations directly from the Master Equation of channel state dynamics and the concentration dynamics of the cytosol and ER. These approaches average Master Equations on (sub)cellular level. This study suggests a theoretical concept not requiring the deterministic limit. So, are there actually differences in the predictions of early mean field theory to our approach and which conceptual distinctions cause them? We would like to discuss a few examples for differences here.

#### The origin of time scales

The representation of the ISI time scale in deterministic models requires necessarily a process realizing that time scale [26, 83, 70]. Slow recovery from negative feedback is one example. Spontaneous spiking behavior of Astrocytes illustrates that this time scale may arise in stochastic models without a slow process. As we know from the measured value of the slope of the moment relation *α* close to 1 (Fig. 4D), the time scale of recovery from negative feedback in Astrocytes is much shorter than the average interspike interval (Fig. 4D, Table 1), and the cell reaches a stationary state soon after a spike. Noise generates the next spike out of this state. If the spike generation probability is small, it takes long before the next spike happens. This long time scale arises from the small value of the spike generation probability. It does not arise from a *slow process* on cell level. When we derive mean field equations for such a system, they do not exhibit spiking and the noise generated time scale is lost. Fitting time constants of putative slow processes to the average ISI leads to erroneous conclusions on mechanisms in these cases. Mean field approaches are not able to capture slow time scales in the generation of individual spikes arising from small probabilities.

#### The relation of rates and time scales to system parameters and variables

Cases like the Astrocytes are identified by *α* ≈1. The case *α* < 1 means spikes occur during the recovery from negative feedback. Since noise strongly affects the spike initiation during the transient, the ISI time scale is not proportional to *λ*^*−*1^ but depends e.g. for the asymptotically symmetric random walk on *λ* like *λ*^*−ν*^ with *ν* varying from 1/2 to close to 1 when *N*_*t*_ changes from 1 to large values [29]. This dependency on the cell variability parameter *N*_*t*_ indicates a potential cell variability of the dependency of the ISI time scale on the relaxation rate.

Rates in mean field equations are proportional to (products of) concentrations according to the law of mass action. Eqs. 3-6 and Figs. 3, 6 show that rates of stochastic processes are proportional to numbers. Thus, mean field approaches predict spike frequencies to be independent of cell size, stochastic approaches can easily explain a dependency on cell size. Mean field equations predict the onset of spiking as bifurcations generating limit cycles, which entail canonical behavior of the period of spiking close to onset. The stimulation response relation does not correspond to any of these canonical relations [37, 36, 55, 70]. The strength of spatial coupling drops out of mean field equations spatially averaging on cell level. However, we see again from Eqs. 3-6, Figs. 3, 6 and the experimental results in Skupin et al. [64] that strength of spatial coupling affects time scales sensitively. While we could imagine mean field approaches taking spatial coupling strength by some effective approximations into account, stochastic approaches have the advantage of working with clusters as discrete entities and thus can naturally accommodate for coupling strength parameters.

### 3.2 Our concept as basis for a comprehensive theory

Our concept not only explains the basic observations, but also defines how to include more detail. The fact that we needed to vary only *λ* and T_min_ to capture the ISI statistics of 4 different pathways illustrates the potential for detail which can be included. For example, more detail on [IP_3_]([A]) could be included by varying *k*_*p*_, [IP_3_]_0_, *g*_0_ and *n*_*r*_. Cell variability in expression levels of pathway components determining [IP_3_]([A]) appears likely to us. Addressing it in the model would require some experimental data to which we could fit such modeling efforts. The parameter 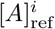 might then turn out to be redundant to other parameters, determining [IP_3_]. Alternative descriptions of CICR and cluster structure can be included by the choice of the CICR-factor in (6) and its dependency on the number of open clusters or channels.

We focused here on puff properties and interspike intervals, since they are sufficient to present the essence of our theory, and the general observations 1-8 refer to them. The transition rates (3), (6) and the Master Equation (7) define a random walk in the state space of the cellular cluster array, which is the mathematical structure of IP_3_ induced Ca^2+^ signaling. They also define the starting point for a more comprehensive theory. The analytical solution for the moments is not limited to the first passage problem for which we used it. Refining *I*-dynamics (replacing the simple linear dynamics (5) we have chosen here) would be the means to include more detailed characteristics of recovery from negative feedback after spike termination and to describe complex signal patterns like bursting [41, 23, 61]. Including processes during the spike in simulations of the model to address spike shape and bursting is straightforward. Hence, we think our approach is able to accommodate the full complexity of Ca^2+^ signals [28, 26].

## Acknowledgments

VNF has been supported by DFG grant Fa350/13-1 to MF.

## Footnotes

1 https://parkerlab.bio.uci.edu/images_movies_presentations/calcium.htm

