## Supplemental Material for "Modeling IP_3_ induced Ca^2+^ signaling based on its interspike interval statistics"


December 20, 2022

#### S1 Calculating moments of the first passage time distribution

An analytic solution for the moments of the first passage time distribution with $n_r$=1 in form of a sum of matrix products has been worked out in Falcke and Friedhoff [1]. We use this solution here and start with the Master equation for the state probabilities $P_k$

$$\frac{dP_k}{dt} = \Psi_{k-1,k}P_{k-1} + (k+1)\delta P_{k+1} - (\Psi_{k,k+1} + k\delta)\, P_k, \quad k = 0, \ldots, N_t - 1. \tag{S1}$$

The first passage time ($t_f$) distribution $F_{0,N_t}(t_f)$ is given by the probability flux out of the state range from 0 to $N_t - 1$:

$$F_{0,N_t}(t_f) = -\frac{d}{dt}\sum_{k=0}^{N_t-1} P_k(t)\Bigg|_{t=t_f} = \Psi_{N_t-1,N_t}P_{N_t-1}. \tag{S2}$$

That equation applies, if we solved the problem with $N_t$ being an absorbing state, i.e. $\Psi_{N_t,N_t-1} = 0$. In matrix notation with the vector of probabilities $P$, we have

$$\frac{dP}{dt} = EP + e^{-\lambda t}DP, \tag{S3}$$

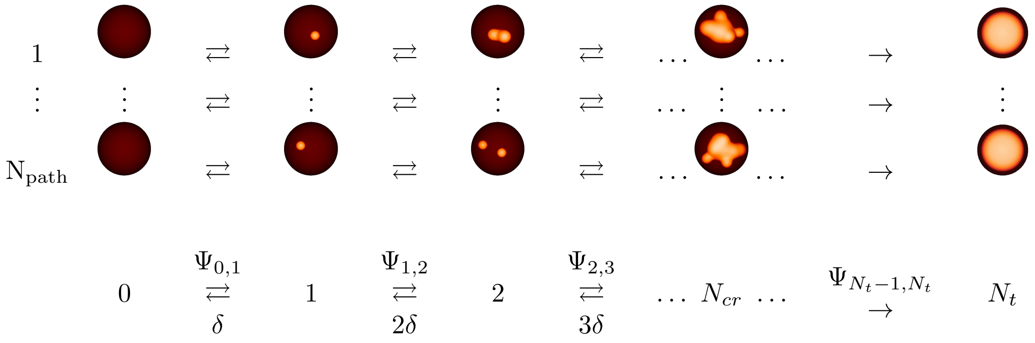


Figure S1: (Top) Open clusters are visualized as small orange spheres. A spike occurs when (almost) all $N_t$ clusters are open. There are $\mathrm{N_{path}}$ paths of cluster openings and closing from 0 to $N_t$ open clusters. (Bottom) Averaging over all paths leads to a state scheme indexed by the number of open clusters.

with the matrices $E$ and $D$ defined by

$$\begin{array}{lcll}\Psi_{k,k+1} & = & g(N_t-k)\left(1+s_pk\right)^n\left(1-e^{-\lambda t}\right), & \\ \psi_{k,k+1} & = & g(N_t-k)\left(1+s_pk\right)^n, & \\ E_{k,k-1} & = & \psi_{k-1,k}, & k=1,\ldots,N_t, \\ E_{k-1,k} & = & k\delta, & k=0,\ldots,N_t-1, \\ E_{k,k} & = & -\left(\psi_{k,k+1}+\delta\right), & k=0,\ldots,N_t-1, \\ D_{k,k-1} & = & -\psi_{k-1,k}, & k=1,\ldots,N_t, \\ D_{k,k} & = & \psi_{k,k+1}, & k=0,\ldots,N_t-1,\end{array} \tag{S4}$$

and all other matrix elements vanish.

The initial condition defines the vector $r$ with $r_0=1$, $r_k=0$, $k=1,\ldots,N_t-1$. The Laplace transform of the Master equation allows for a comfortable calculation of moments of the first-passage times. The Laplace transform of (S3) is the system of linear difference equations [1]

$$s\tilde{P}(s)-r_i \quad = \quad E\tilde{P}(s)+D\tilde{P}(s+\lambda). \tag{S5}$$

We define $\tilde{B}(s)=\left(\mathbb{1}s-E\right)^{-1}D$ and obtain [1]

$$\tilde{P}(s) \quad = \quad \left(\mathbb{1}s-E\right)^{-1}r+\sum_{k=1}^{\infty}\prod_{j=0}^{k-1}\tilde{B}(s+j\lambda)\left[\mathbb{1}(s+k\lambda)-E\right]^{-1}r \tag{S6}$$

as the solution of (S5). The Laplace transform of the first-passage-time distribution is [1]

$$\tilde{F}_{0,N_t}(s) \quad = \quad \psi_{N_t-1,N_t}\left[\tilde{P}_{N_t-1}(s)-\tilde{P}_{N_t-1}(s+\lambda)\right], \tag{S7}$$

The moments of the first-passage-time distribution are given by [5]

$$\langle {t_f}^n\rangle=(-1)^n\frac{\partial^n}{\partial s^n}\tilde{F}_{0,N_t}(s)\bigg|_{s=0}. \tag{S8}$$

### S2 Numerical methods

#### S2.1 Evaluation of Eq. S6

If $N$ is large and the ratio between the fastest state transition rate and the relaxation rate $\gamma$ is large, high precision of the numerical calculations is required for the use of (S6). *A priori* known values like $\tilde{F}_{0,N_t}(s=0)=1$ can be used to monitor the convergence of the calculations. The matrix products become very large at intermediate values of $j\lambda$ during the summation in (S6), and their sign alternates such that two consecutive summands nearly cancel. Intermediate summands are of order larger than $10^{17}$, and thus we face loss of significant figures even with the numerical floating-point number format long double. We used Arb, a C numerical library for arbitrary-precision interval arithmetic [2], to circumvent this problem. It allows for arbitrary precision in calculations with (S6). Computational speed is the only limitation with this library and has determined the parameter range for which we established analytical results. We were able to go to a time-scale separation of $\approx 10^{-6}$ using this library. We calculated moments with a relative precision of 4 $10^{-4}$. Arbitrary precision computations with Arb take between 1 s and 10 days, heavily dependent on the matrix dimensions set by the index of the absorbing state. See Falcke and Friedhoff [1] for more detail.

#### S2.2 Simulation method

To be able to also compute first passage times for $n_r \neq 1$ and to allow for sampling of ISIs for better comparison of results matching experimental workflows, we also simulated the state chain shown in Fig. S1 stochastically using a Gillespie algorithm in C++. We use a discrete chain of states to model the number of open clusters $k$ from $0\leq k\leq N_t$ in discrete time steps on the order of $dt=10^{-5}$ $s$. The up- and down-rates from our model are given by

$$\Psi_{k,k+1} \quad = \quad g\ (N_t-k)\left(1+s_pk\right)^n\left(1-e^{-\lambda t}\right)^{n_r} \tag{S9}$$

$$\Psi_{k,k-1} \quad = \quad k\ \delta \tag{S10}$$

and are used to determine if the state variable $k$ stays constant or changes in the current time window of $dt$ at time $t$ and, in case of a change, if the jump occurs up or down. If a state $k$ is reached for the first time at time $t = t_k$, the first passage time $t_k = F_i(k)$ is stored for evaluation. After reaching state $N_t$, or typically $k = 20$ because typically $N_{cr} < 10$, the simulation is considered finished (a spike has occurred), and another iteration is initiated. The first passage times $F_i(k)$ of iteration $i$ for all states $k$ are considered a sample of the probability mass function of the underlying stochastic process and are saved to a file. Typically, around $10^4$ iterations were used to compute the approximation of the probability mass function for one set of parameters $(N_t, s_p, \lambda, n_r)$ with $\delta$ and $n$ being constant for all simulations. To compute $F_i(g)$ for one set of parameters, g was varied inhomogeneously between $10^{-3}\ s^{-1}$ and $40\ s^{-1}$ (Fig. S3A). To reach $F(g)$, we averaged over iterations, $F(g) = \langle F_i(g) \rangle$.

For small values of g for some parameter sets $F_i(g)$ would sometimes not converge, i.e. the stochastic process would not reach state $k = N_t$ and would idle around $k = 0$ for a long time. To overcome this, we introduced a cutoff at 4000 s, after which the current iteration was terminated and the next one initiated. If there was a low ratio of successful runs to initiated ones, the simulation for that case was canceled all together. Due to the cutoff affecting the quality of the approximation of the probability mass function of the underlying stochastic process, precision of the resulting $F(g)$ was assumed only to hold for $F(g) < 10^3\ s$, i.e. larger results were discarded from evaluation. This agrees with data from experiments that we used as input for our model, where generally $\mathrm{T_{av}}$ ¡ 600 s holds. Agreement of analytical computations using the method from Falcke and Friedhoff [1] with simulations is very good, Fig. (S2).

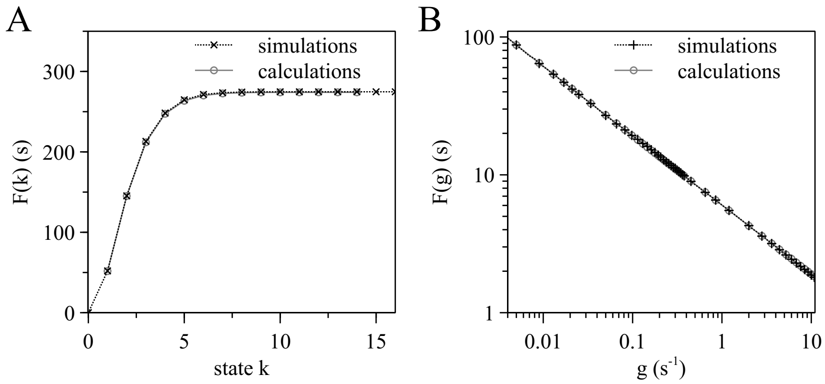


Figure S2: Comparison of analytical (grey, circles, solid) and simulation (black, crosses, dotted) results of the average first passage time, here both in $F(k)$ and $F(g)$, shows good agreement. **(A)** $F(k) = \langle F_i(k) \rangle$ for simulations and $F(k) = -\partial \tilde{F}_{0,k}(s)/\partial s$ at $s = 0$ (Eq. S8) for analytical results with parameters $\lambda = 0.001443\ s^{-1}$, $s_p = 1.70$, $N_t = 20$ and $g = 0.021\ s^{-1}$. Analytical calculations were computed for $1 \leq k \leq 14$, while simulations ran for $1 \leq k \leq 20$. Here $N_{cr} = 7$ sets the convergence behavior, i.e. $F(k = N_{cr}) \approx F(k > N_{cr})$. $F(k)$ converges against $F(N \geq N_{cr}) \approx 274.1\ s$ for calculations and $F(N \geq N_{cr}) \approx 274.7\ s$ for 11k simulations. We set $\mathrm{T_{av}} = F(k = N_t - 2) = F(k = 18)$. **(B)** Double-log plot showing F(g) for $\lambda = 0.001443\ s^{-1}$, $s_p = 1.70$, $N_t = 30$, and $N = 1$. Since $N = 1$ holds, we only used the analytical solution from eq. (5) in [3]. 11k to 15k simulations per data point were used for averages.

### S3 Determining parameters of the model from experiments

We describe in the following how we determine parameter values for a given cell type and agonist. The input values are the range of $\mathrm{T_{av}}$ values, the slope of the moment relation $\alpha$, and the stimulation response relation $\mathrm{T_{av}}([\mathrm{A}])$ .

The solution of the first passage problem provides the average first passage time $F(g, s_p, N_t, \lambda, n_r)$ in dependence on the parameters among them the puff rate $g$. The first passage time plus a constant equals the average interspike interval $\mathrm{T_{av}}$. The known dependencies of the population average of $\mathrm{T_{av}}$ on the agonist concentrations for 4 different pathways reported in [4] provides the opportunity to determine the relation between the puff rate $g$ and the agonist concentration $A$. We start with

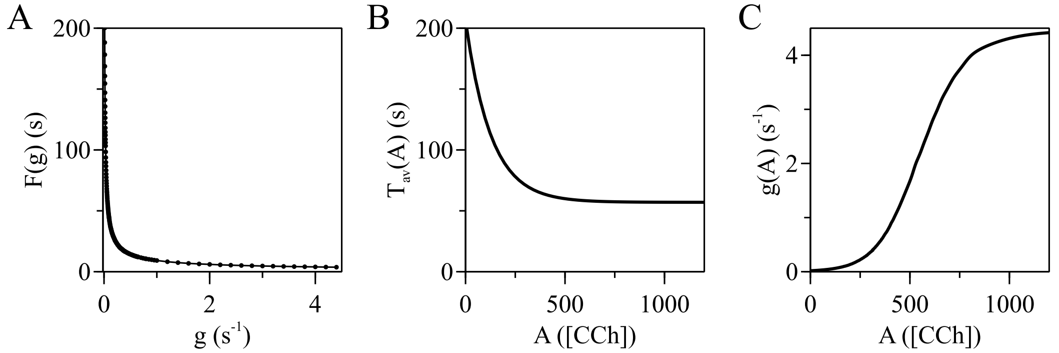


Figure S3: **(A)** Data points are results of stochastic process $F^r(g)$ for reference case of HEK-CCh pathway, $N_t = 30$ and $s_p = 2.2$. Solid line is a spline. $F^i(g)$ converge to 0 for large $g$. **(B)** Stimulation response relation $\mathrm{T_{av}(A)}$ for HEK-CCh pathway with values from experiment, $\mathrm{T_{min}} = 57\ s$, $\gamma = 0.00784\ \mu M^{-1}$, $[A]^i_{ref} = 30\ \mu M$, and $\mathrm{T_{ref}} = 175$ s [4]. Here we plotted it also for $[A] < [A]^i_{ref}$. $\mathrm{T_{av}(A)}$ converges against $\mathrm{T_{min}}$ for large [A]. **(C)** g([A]) computed via eq. (S14). We used $[A]' = 1200\ \mu$M and $g' = 4.5\ s^{-1}$, resulting in $\Theta \approx -3.681\ s$.

identifying a range of parameter values for $s_p$, $N_t$, $\lambda$ and $n_r$ providing the range of $\mathrm{T_{av}}$ values and the slope $\alpha$ observed in experiments. Since the value of $\lambda$ and $n_r$ are the same for all cells of a given type and pathway, we drop them from the list of parameters in the following.

#### S3.1 From stochastic process to stimulation response relation

To map the results of the stochastic process, i.e., the first passage time $F(g, s_p^i, N_t^i) = F^i(g)$ onto the stimulation response relation

$$T_{av}([A]) = (T^i_{ref} - T_{\min})\ e^{-\gamma\left([A]-[A]^i_{ref}\right)} + T_{\min} \tag{S11}$$

from experiment, we will apply a non-linear coordinate transformation g([A]). It will be determined for the reference case and then used for all other cases as well. Our ansatz consists of adding two constants $\mathrm{T_{min}}$ and $\Theta$ to the first passage time $F$. $\Theta$ accounts for the contribution of $F(g, s_p^i, N_t^i)$ to $\mathrm{T_{min}}$ :

$$F^i(g) + \Theta_i + T_{\min} = (T^i_{ref} - T_{\min})\ e^{-\gamma\left([A]-[A]^i_{ref}\right)} + T_{\min} \tag{S12}$$

For the reference case $i = r$ with $N_t^r$ and $s_p^r$ we find

$$\Theta_r(g', [A]') = (T^r_{ref} - T_{\min})\ e^{-\gamma\left([A]'-[A]^r_{ref}\right)} - F^r(g'). \tag{S13}$$

The values of $g'$ and $[A]'$ allow us to fix $g([A])$ at the point $g([A]') = g'$. This degree of freedom corresponds to choosing which part of $F^i(g)$ is mapped onto $\mathrm{T_{av}(A)}$ by setting the value of $\Theta_r$. We generally choose large values for $[A]'$, even outside the experimental measured range, to set the saturation value of $g([A])$ to $g'$.

We can now determine $g([A])$ by using the inverse of the stochastic process $F(g)$, i.e. $g = F^{-1}(F(g))$, where we express $F(g)$ via $\mathrm{T_{av}(A)}$ applying eq. (S12). Using the reference case, we find

$$\begin{aligned} g^r([A]) &= F_r^{-1}\left(T^r_{\mathrm{av}}([A]) - \Theta_r - T_{\min}\right) \\ &= F_r^{-1}\left(\left(T^r_{ref} - T_{\min}\right)\left(e^{-\gamma([A]-[A]^r_{ref})} - e^{-\gamma([A]'-[A]^r_{ref})}\right) + F_r(g')\right) \end{aligned} \tag{S14}$$

$\Theta_r$ and $g^r([A])$ specify together with $\lambda$ the cell type and agonist. With the expression for $g^r([A])$ (Eq. S14) and $\Theta_r$ (Eq. S13), and applying them also to the non-reference cases, we reach

$$T_{\mathrm{av}}([A], s_p^i, N_t^i) = \begin{cases} = F(g^r([A]), s_p^i, N_t^i) + \Theta_r + T_{\min} & , [A] \geq [A]^i_{ref} \\ 0 & , [A] < [A]^i_{ref} \end{cases}, \tag{S15}$$

where $s_p^i$, $N_t^i$, $[\mathrm{A}]^i_{ref}$ are the parameters specifying individual cells.

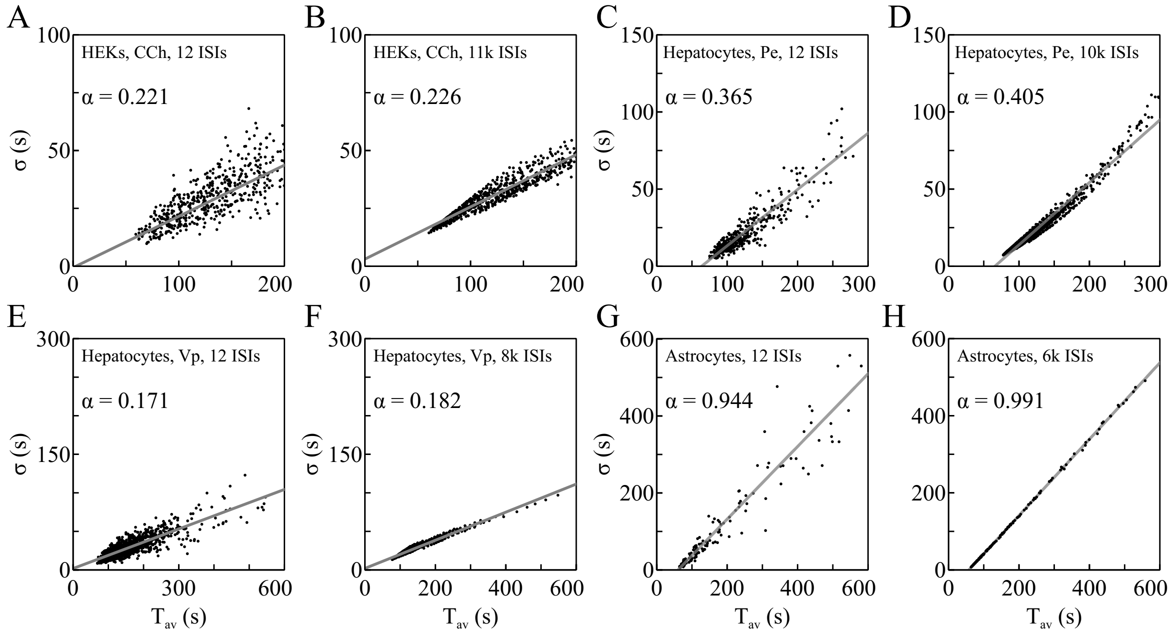


Figure S4: $\sigma$-$\mathrm{T_{av}}$ plots showing slightly varying $\alpha$ for four pathways in case of sampling the results with 12 ISIs (A, C, E, G) vs using a very larger number of ISIs ¿ 6k (B, D, F, H).

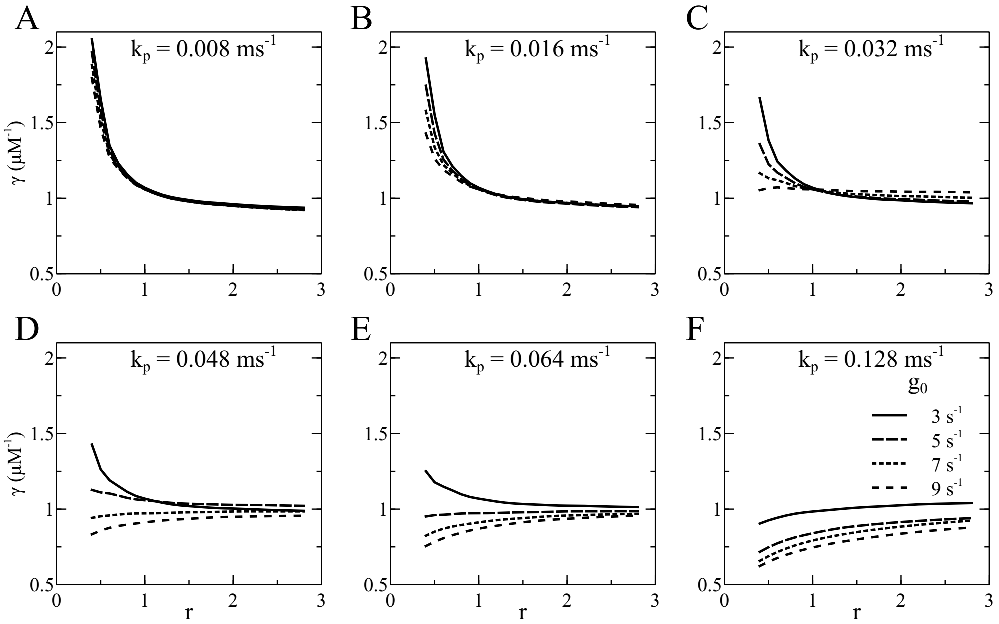


Figure S5: The relation between the agonist sensitivity $\gamma$ of the stimulation response relation Eq. S11 and the saturation concentration of the $IP_3$ pathway $[IP_3]_{\mathrm{max}}$. The ratio of the reference value $[IP_3]_{\mathrm{ref,max}}$ to $[IP_3]_{\mathrm{max}}$ is denoted r: r=$[IP_3]_{\mathrm{max}}/[IP_3]_{\mathrm{ref,max}}$. The reference value of $\gamma$ is 1.059 $\mu$M$^{-1}$. With each pair of $k_p$ and $g_0$, we varied $[IP_3]_{\mathrm{max}}$ in the calculation of $T_{\mathrm{av}}(g([IP_3]([A])))$ and fitted the result to an exponential function to determine $\gamma$. The $[IP_3]$ sensitivity of Eq. 8 is $k_p$, and $g_0$ is the maximum puff rate. There is a range of r-values where $\gamma$ is essentially constant for each pair of $k_p$ and $g_0$.

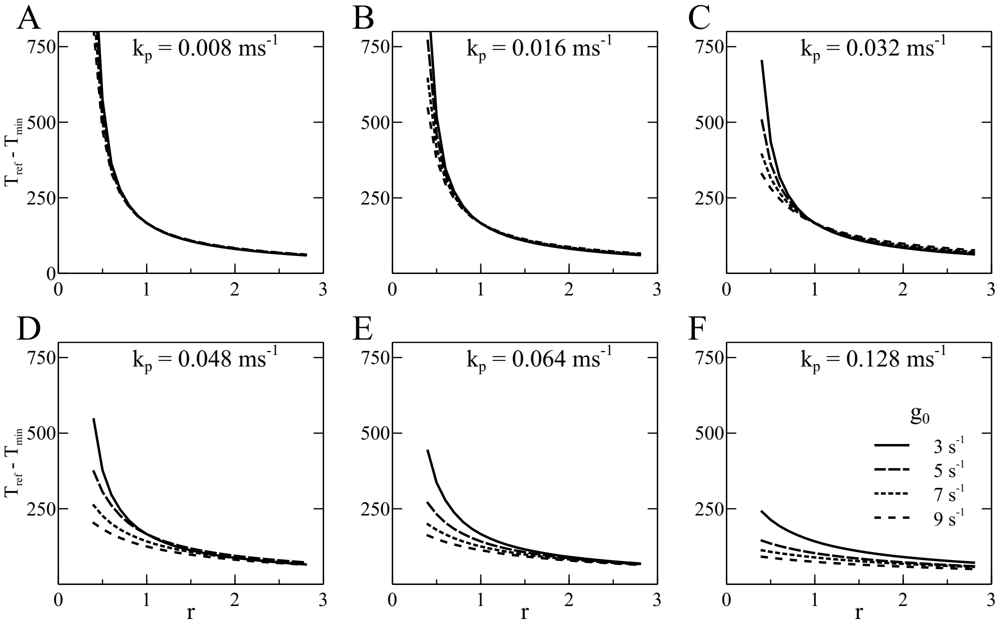


Figure S6: $T_{ref} - T_{min}$ against ratio $r$ for different $k_g$ and $g_0$.

[5] N.G. van Kampen. *Stochastic processes in physics and chemistry.* North-Holland, Amsterdam, 2001.
